# Clinical *Candida–Lactobacillus* Isolate Pairs Reveal Distinct Host-Pathogen Interactions in a Vaginal Epithelial Model

**DOI:** 10.1101/2025.09.24.678373

**Authors:** Esraa Mubarak Almutawa, Lily Novak-Frazer, Raveen Tank, Elaine M. Bignell, Riina Rautemaa-Richardson

**Author notes:** **Funding** Esraa Mubarak Almutawa was funded by the Public Authority for Applied Education and Training (PAAET), Kuwait.

## Abstract

Vulvovaginal candidiasis (VVC) is a highly prevalent condition affecting up to 75% of women, yet the mechanisms underpinning the transition from asymptomatic colonisation by Candida albicans to symptomatic infection remain incompletely understood. VVC is hormonally driven and rare in young girls and postmenopausal women. A central challenge lies in dissecting the complex interplay between microbial communities, hormonal factors, and host epithelial responses. While C. albicans is part of the normal vaginal microbiota at low abundance, its morphogenic switch to the hyphal form is closely linked with pathogenicity. Concomitantly, the dominance of specific Lactobacillus species is known to modulate vaginal health, but their precise role in shaping host-pathogen interactions during VVC is unresolved.

We hypothesised that clinical isolate pairs of C. albicans and Lactobacillus spp. may differ in their ability to drive host epithelial damage, and that hormonal signals such as estrogen may exacerbate these interactions. To test this, we developed an in vitro VVC model using human vaginal epithelial cell (VEC) monolayers co-cultured with clinical isolate pairs. Using LDH release as a marker of cytotoxicity, extracellular pH monitoring, and immunoassays of mitogen-activated protein kinase (MAPK) phosphorylation, we interrogated host responses across microbial conditions in the presence or absence of estrogen.

Our findings demonstrate that two clinical isolate pairs exhibit divergent behaviours. Pair C (C. albicans 203 and L. gasseri 167) suppressed epithelial cytotoxicity and maintained acidic pH, while Pair D (C. albicans 205 and L. jensenii 172) enhanced host damage and did not acidify the environment. Morphological differences in C. albicans isolates correlated with MAPK activation profiles, supporting a phenotype-specific host response. These results reveal that strain-level interactions within the vaginal microbiota significantly modulate host-pathogen dynamics.

## Introduction

Vulvovaginal candidiasis (VVC) is a highly prevalent condition affecting up to 75% of women, yet the mechanisms underpinning the transition from asymptomatic colonisation by *Candida albicans* to symptomatic infection remain incompletely understood. VVC is hormonally driven and rare in young girls and postmenopausal women. A central challenge lies in dissecting the complex interplay between microbial communities, hormonal factors, and host epithelial responses. While *C. albicans* is part of the normal vaginal microbiota at low abundance, its morphogenic switch to the hyphal form is closely linked with pathogenicity^1^. Concomitantly, the dominance of specific *Lactobacillus* species is known to modulate vaginal health, but their precise role in shaping host-pathogen interactions during VVC is unresolved.

We hypothesised that clinical isolate pairs of *C. albicans* and *Lactobacillus* spp. may differ in their ability to drive host epithelial damage, and that hormonal signals such as estrogen may exacerbate these interactions. To test this, we developed an *in vitro* VVC model using human vaginal epithelial cell (VEC) monolayers co-cultured with clinical isolate pairs. Using LDH release as a marker of cytotoxicity, extracellular pH monitoring, and immunoassays of mitogen-activated protein kinase (MAPK) phosphorylation, we interrogated host responses across microbial conditions in the presence or absence of estrogen.

Our findings demonstrate that two clinical isolate pairs exhibit divergent behaviours. Pair C (*C. albicans* 203 and *L. gasseri* 167) suppressed epithelial cytotoxicity and maintained acidic pH, while Pair D (*C. albicans* 205 and *L. jensenii* 172) enhanced host damage and did not acidify the environment. Morphological differences in *C. albicans* isolates correlated with MAPK activation profiles, supporting a phenotype-specific host response^2^. These results reveal that strain-level interactions within the vaginal microbiota significantly modulate host-pathogen dynamics.

## Results

### *Candida albicans* clinical isolates differ in morphology and epithelial cytotoxicity

To establish a VVC-relevant *in vitro* system, we tested clinical *C. albicans* isolates (203 and 205) for their ability to induce epithelial cell damage. Both isolates were applied at a standardised load (10^5 cells/ml) to VEC monolayers. *C. albicans* 203 exhibited extensive hyphal formation, while *C. albicans* 205 maintained a predominantly yeast morphology. This morphogenetic divergence was associated with increased cytotoxicity, as measured by lactate dehydrogenase (LDH) release (Figure 1).

### MAPK activation is isolate- and morphology-dependent

We next analysed the host signalling response using immunoassays to measure MAPK phosphorylation. VECs exposed to *C. albicans* 203 showed significantly higher activation of ERK1/2 and p38 at 4- and 8-hours post-infection, correlating with enhanced hyphal switching, compared to *C. albicans* 205. JNK activation was not significantly different between isolates, suggesting ERK1/2 and p38 are key pathways discriminating virulent morphotypes (Figure 2).

**Figure 1.**
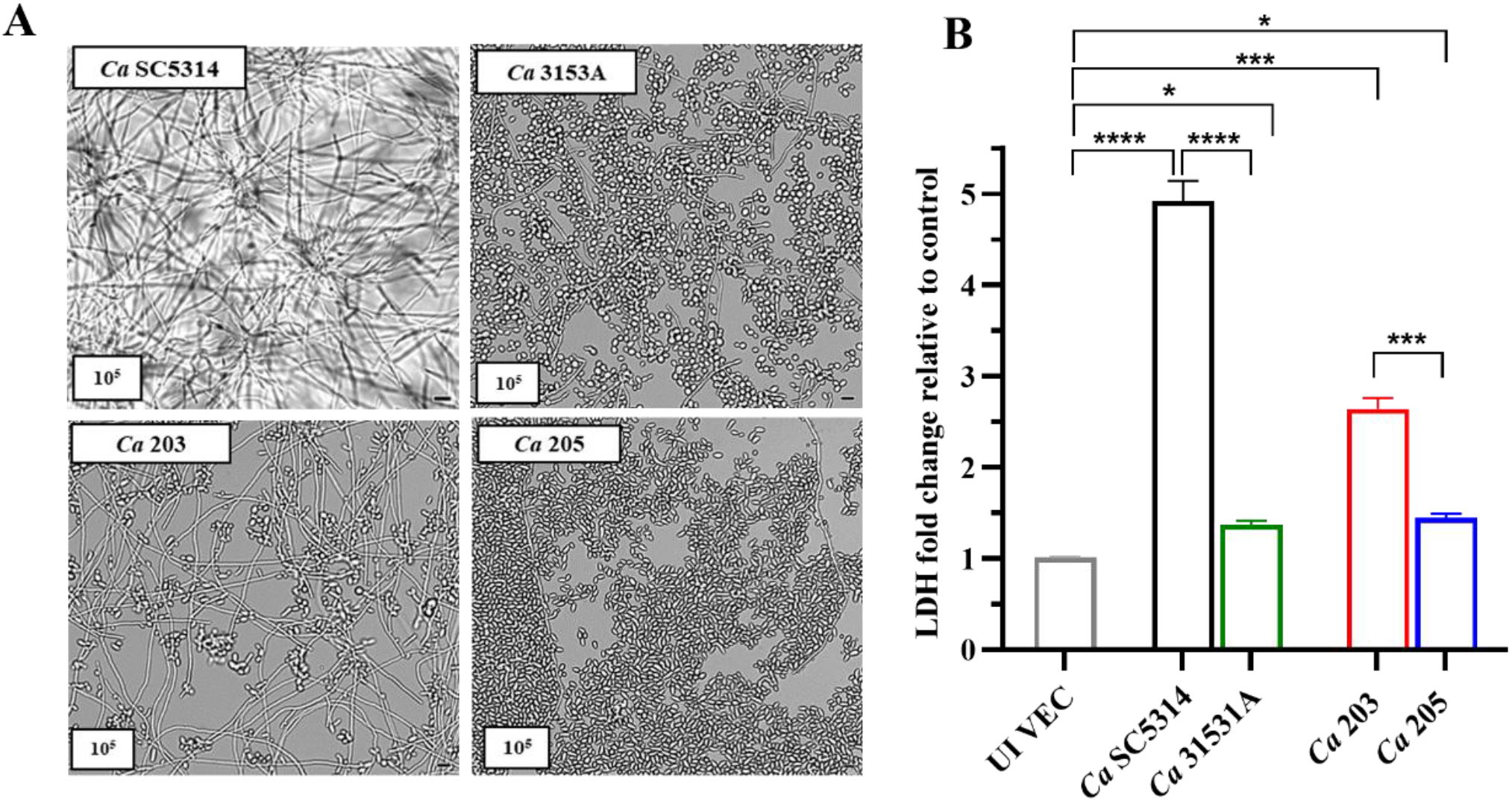
Morphology and cytotoxicity of *C. albicans* clinical isolates. (A) Representative micrographs showing yeast vs hyphal forms. (B) LDH release in VEC monolayers exposed to *C. albicans* 203 or 205 (mean ± SEM, n = 3).

**Figure 2.**
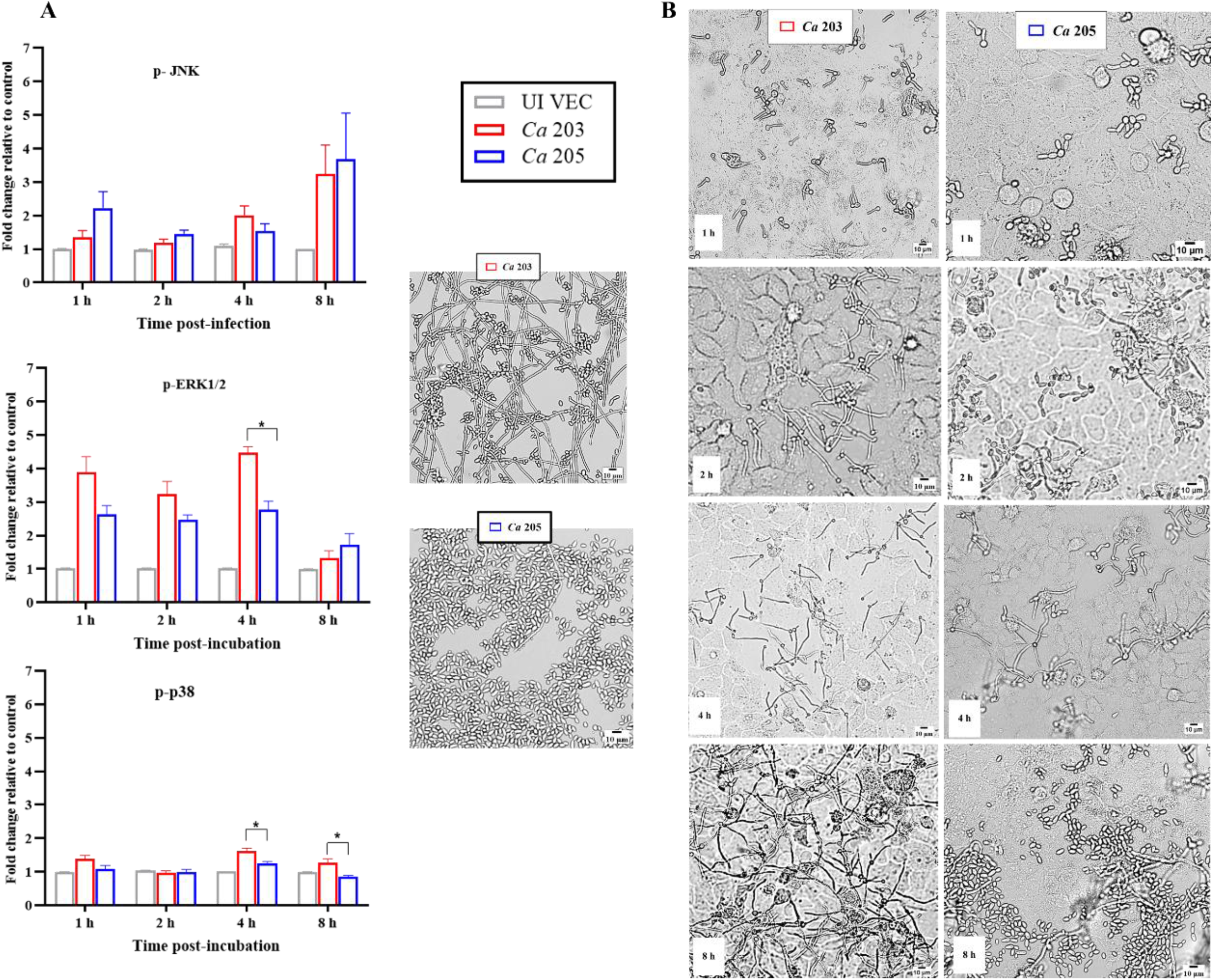
MAPK signalling activation in response to *C. albicans* (*Ca*) clinical isolates. (A) Fold change in JNK, ERK1/2, and p38 phosphorylation following infection with *Ca* 203 or *Ca* 205. (B) Time-course of MAPK activation over 8 h.

### *Lactobacillus* clinical isolates modulate host damage in a species-specific manner

To determine the influence of bacterial co-residents, we co-cultured VECs with matched clinical pairs: Pair C (*C. albicans* 203 with *L. gasseri* 167) and Pair D (*C. albicans* 205 with *L. jensenii* 172). *L. gasseri* suppressed LDH release and acidified the extracellular environment (pH ≤ 4.3), while *L. jensenii* failed to suppress cytotoxicity and did not significantly alter pH (Figures 3 and 4). Thus, protective versus non-protective behaviour of *Lactobacillus* spp. may reflect functional heterogeneity at the species or strain level.

**Figure 3.**
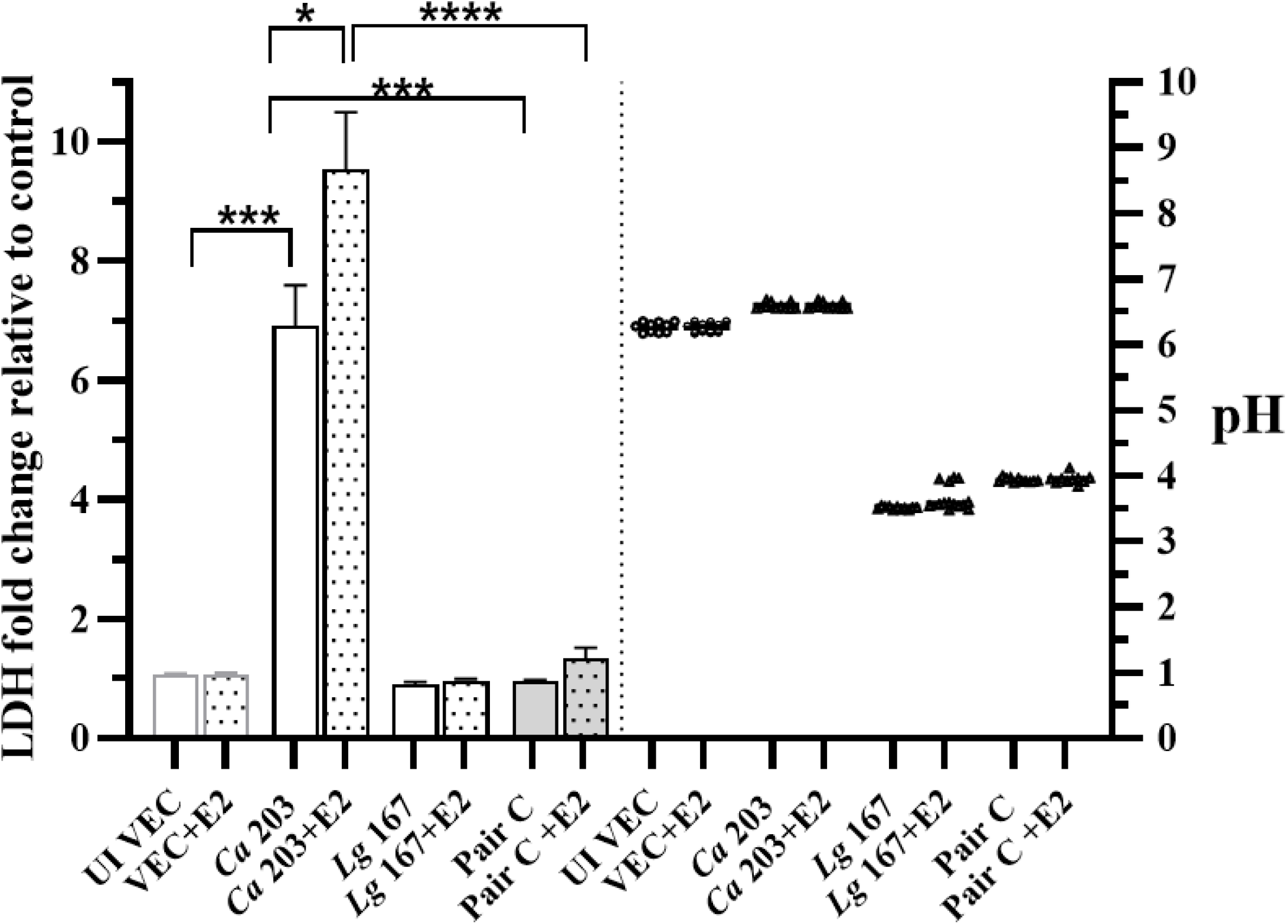
Host cytotoxicity and pH modulation by *L. gasseri* (*Lg*) 167 and estrogen effect on *C. albicans* 203-induced damage. LDH release from VECs co-cultured with Pair C (*Ca* 203 and *Lg* 167) with or without 17β-estradiol (E2) (1 nM) (left axis). Extracellular pH measurements across conditions (right axis). pH remained stable across estrogen treatments.

**Figure 4.**
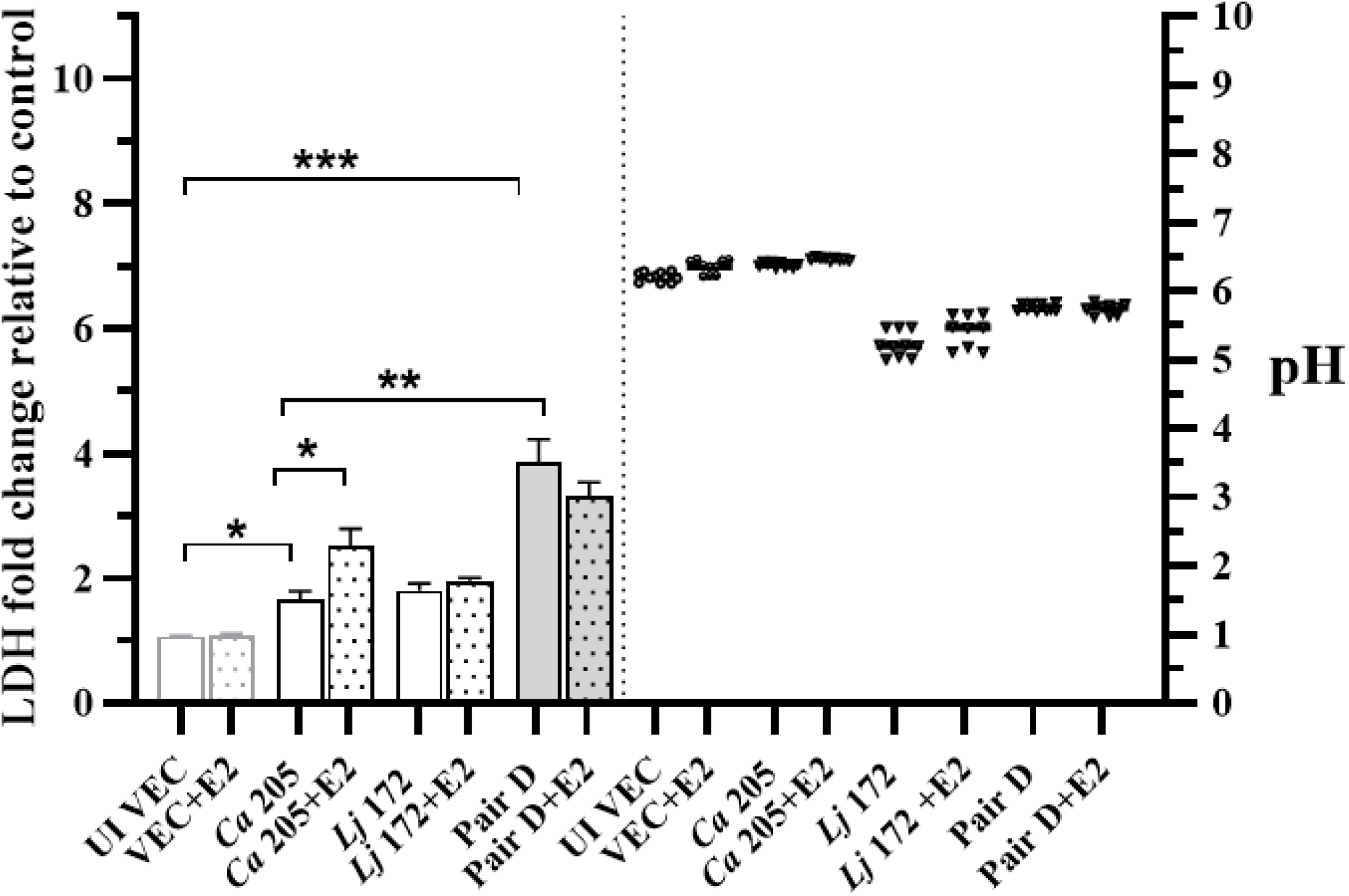
Host cytotoxicity and pH modulation by *L. jensenii* (*Lj)*172 and estrogen effect of on *C. albicans* 205-induced damage. LDH release from VECs co-cultured with Pair D, comprising *Ca* 205 and *Lj* 172, with or without 17β-estradiol (E2) (1 nM) (left axis). Extracellular pH measurements across conditions (right axis). pH remains stable across estrogen treatments.

### Estrogen enhances *C. albicans*-induced cytotoxicity independent of pH

Estrogen (17β-estradiol) significantly enhanced epithelial damage caused by both *C. albicans* isolates. Notably, this effect was independent of extracellular pH, suggesting a direct enhancement of fungal virulence or altered host susceptibility (Figures 3 and 4). Estrogen alone did not induce VEC cytotoxicity and monolayer integrity was unaffected.

Together, these data reveal that the interaction between *C. albicans* and *Lactobacillus* clinical isolates can be either protective or pathogenic, depending on the strain pairing. Our model provides a tractable system for dissecting host-microbiota-pathogen interactions and underscores the need for strain-level resolution in studying vaginal health and disease.

## Discussion

This study presents a refined *in vitro* model for interrogating multifactorial interactions underlying VVC pathogenesis, emphasising the critical role of strain-level microbial behaviour. We demonstrate that clinical isolate pairs of *C. albicans* and *Lactobacillus* spp. display contrasting impacts on host epithelial integrity. Specifically, *C. albicans* 203—exhibiting extensive hyphal growth—elicited higher cytotoxicity and stronger MAPK activation than *C. albicans* 205, which retained a predominantly yeast morphology^2^. These morphogenetic traits aligned with activation of host ERK1/2 and p38 pathways, supporting their role in epithelial discrimination of pathogenic fungal forms^1^.

Importantly, *Lactobacillus* co-culture outcomes were isolate-specific. *L. gasseri* 167 suppressed *C. albicans*-induced epithelial damage and lowered extracellular pH, in contrast to *L. jensenii* 172, which failed to confer protection.

The inclusion of estrogen further revealed that host susceptibility to fungal cytotoxicity is hormone-sensitive^4^, even when pH remains unaltered. This aligns with epidemiological observations linking elevated estrogen levels with increased VVC prevalence^4^, especially in reproductive-age women. Recent findings indicate that estrogen also enhances *C. albicans* virulence and impairs innate immune responses^3^, further supporting its role in VVC pathogenesis.

Our findings argue for a precision microbiome approach to vaginal health, where both fungal and bacterial strain identity profoundly shape disease trajectories. Future therapeutic strategies may benefit from strain-specific diagnostics and the targeted use of probiotics such as *L. gasseri*. Moreover, this platform allows exploration of additional host-pathogen variables—such as glucose levels, immune mediators, and microbiome-derived metabolites—that may influence VVC onset or recurrence.

## Materials and Methods

### Microbial strains and epithelial cell culture

Clinical isolates of *Candida albicans* (203 and 205) and co-isolated *Lactobacillus* spp. (*L. gasseri* 167 and *L. jensenii* 172) were sourced from patients with VVC. *C. albicans* reference strains SC5314 and 3153A were used for baseline comparisons. All yeasts were grown aerobically on Sabouraud dextrose agar; *Lactobacillus* strains were cultured anaerobically on MRS agar. Vaginal epithelial cells (A431 cell line) were maintained in complete DMEM with 10% FBS and 1% penicillin-streptomycin. Cells were seeded into 24-well or 6-well plates to reach ∼90% confluence prior to infection.

### Infection assays and co-culture model

Standardised microbial inocula (10^5 cells/ml *C. albicans* and 10^7 CFU/ml *Lactobacillus*) were applied individually or in combination to serum-starved VEC monolayers. Infections were conducted for up to 24 hours at 37°C, 5% CO2. Estrogen (17β-estradiol) was applied at 1 nM where indicated. Extracellular pH was measured post-infection using a microelectrode probe.

### Cytotoxicity and epithelial integrity assays

LDH release in culture supernatants was quantified using a colorimetric LDH assay kit. VEC detachment assays were performed using DAPI staining and automated fluorescence microscopy to assess monolayer integrity. Image analysis was conducted in FIJI.

### MAPK phosphorylation profiling

VEC monolayers were lysed at 1, 2, 4, and 8 h post-infection. Total protein was extracted and normalised to 200 µg/ml for use in multiplex Bio-Plex immunoassays. Phosphorylated ERK1/2, JNK, and p38 levels were measured and expressed as fold change relative to uninfected controls.

### Statistical analysis

All data represent the mean of three independent biological replicates unless otherwise noted. Error bars indicate SEM. Data were analysed using GraphPad Prism v9.0. Statistical significance was determined by one-way ANOVA or unpaired t-test as appropriate (p < 0.05).

### Figures

All data represent the mean of three independent biological replicates unless otherwise noted. Error bars indicate SEM. Statistical significance was determined by one-way ANOVA or unpaired t-test as appropriate.

## Acknowledgements

Esraa Mubarak Almutawa was funded by a government grant from PAAET, Kuwait, and supported by the Kuwait Culture Office, Kuwait Embassy, UK (KCOUK).

R.R.-R. is supported in part by the National Institute for Health and Care Research (NIHR) Manchester Biomedical Research Centre (NIHR203308), UK.

## Notes

### Competing Interest Statement

The authors have declared no competing interest.

